# Boosting Brainpower in Ageing: Task-Specific and Transfer Effects of tDCS-Enhanced Working Memory Training

**DOI:** 10.1101/2025.02.19.639019

**Authors:** Esteban León-Correa, Philipp Ruhnau, Alex Balani, Adam Qureshi, Dorothy Tse, Stergios Makris

## Abstract

Ageing is associated with neural alterations that impair cognitive abilities, particularly working memory (WM)—the capacity to temporarily store and manipulate information essential for daily functioning. Transcranial Direct Current Stimulation (tDCS) has shown promise in enhancing WM in older adults by modulating cortical excitability and promoting neuroplasticity. However, findings on the combined effects of tDCS and WM training remain inconsistent, particularly regarding transfer effects to non-trained WM tasks (near transfer) or other cognitive domains (far transfer).

This study examined the behavioural and neural effects of tDCS on distinct WM phases (encoding, retention/manipulation, recall) in older adults. Over three weeks, participants underwent six sessions of adaptive WM training paired with tDCS targeting the left dorsolateral prefrontal cortex (DLPFC). Neural activity was assessed via EEG, focusing on theta and alpha bands. Participants were divided into tDCS and placebo groups and completed two WM tests, the Letter Span and Corsi Test, with retention and manipulation conditions.

Results showed no additional behavioural or neural benefits of tDCS. However, both groups exhibited near-transfer effects that persisted one-month post-training. Neural changes varied by task: the Letter Span showed increased alpha power during manipulation, indicating enhanced maintenance of information during that phase, while the Corsi Test showed reduced left-hemisphere theta power during recall, reflecting reduced hemispheric asymmetry.

Inter-individual variability likely contributed to inconsistencies and a lack of correlations between outcomes. These findings highlight the importance of personalised tDCS protocols and suggest adaptive WM training can yield enduring cognitive and neural benefits in older adults.

## INTRODUCTION

The global population of individuals aged 60 and over has increased significantly in recent years and is projected to more than double within the next 30 years (UN Department of Economic and Social Affairs - Population Division, 2020). While this demographic shift demonstrates that people are living longer, it also presents numerous medical and social challenges. Ageing is associated with pronounced structural and functional neural alterations that compromise cognitive abilities, leading to a diminished quality of life and loss of independence (Cabeza et al., 2018; D. C. Park & Festini, 2017; Rieck et al., 2017; Sala-Llonch et al., 2015; Salthouse, 2010).

One of the most affected cognitive domains in ageing is working memory (WM; Grady, 2012; Salthouse, 2011; Ziaei et al., 2017). WM refers to the ability to store and manipulate information temporarily to complete a task successfully (Baddeley, 1998, 2003). It is integral to daily functioning, underpinning processes such as decision-making, problem-solving, and fluid intelligence (Baddeley, 2012; Conway et al., 2003; Engle et al., 1999). Transcranial Direct Current Stimulation (tDCS) has emerged as a promising intervention for enhancing WM in older adults, owing to its capacity to modulate cortical excitability and promote neuroplasticity (Chan et al., 2021a; Nitsche et al., 2003; Nitsche & Paulus, 2000; Priori et al., 1998). This non-invasive technique administers weak electrical currents to the scalp through anodal and cathodal electrodes. The anodal electrode increases cortical excitability, while the cathodal electrode has an inhibitory effect (Nitsche et al., 2003; Nitsche & Paulus, 2000; Priori et al., 1998). To improve its efficacy, tDCS is often combined with cognitive training to activate the targeted neural circuits (Giordano et al., 2017; Stagg et al., 2018). Researchers propose that coupling WM training with multiple tDCS sessions may optimise outcomes, with cumulative gains over sessions potentially inducing long-term potentiation (LTP), thereby fostering enduring neuroplastic changes and cognitive improvements (Berryhill & Martin, 2018; Chan et al., 2021a; Monte-Silva et al., 2013). Most studies aiming to enhance WM performance through tDCS have focused on the dorsolateral prefrontal cortex (DLPFC), given its central role in key executive functions, inhibitory control, maintenance and manipulation of information, updating processes, cognitive flexibility, and decision-making (Jimura et al., 2018; C. Kim et al., 2015; Murty et al., 2011; Osaka et al., 2003; Rodriguez Merzagora et al., 2014; Vartanian et al., 2013). Furthermore, the DLPFC operates within a fronto-parietal network, which encompasses other critical regions involved in information processing and memory consolidation. These include the anterior cingulate cortex (ACC), parietal cortex (PC), hippocampus, thalamus, caudate nucleus, default mode network, and cerebellum, among others (Andersen & Cui, 2009; Bolkan et al., 2017; Chein et al., 2011; Gottwald et al., 2004; C. Kim et al., 2015; A. B. Moore et al., 2013; Murty et al., 2011; Nissim et al., 2017; Osaka et al., 2003; Owen et al., 2005; Ziemus et al., 2007).

However, findings from studies investigating the combined effects of tDCS and WM training in older adults remain inconsistent and, in some cases, contradictory (Indahlastari et al., 2021; Siegert et al., 2021; Summers et al., 2016). While most research frequently demonstrates that tDCS enhances learning effects (i.e., performance improvements across training sessions) more significantly than cognitive training alone (Perceval et al., 2016; Satorres et al., 2022), results of transfer of training gains to non-trained WM tasks (near-transfer effects) and other cognitive domains (far-transfer effects) have been mixed. For instance, Park et al. (2014) observed improved verbal WM following ten sessions of tDCS but found no enhancement in visual WM, nor any evidence of transfer to broader cognitive domains such as attention and executive functioning (far-transfer effects). In contrast, Stephens & Berryhill (2016) found no near-transfer effects but noted both training effects and sustained far-transfer effects, including enhanced processing speed, cognitive flexibility, and improvements in everyday functioning that persisted for up to one month post-training. Similarly, Jones et al. (2015) reported that, while both tDCS and control groups benefitted from the intervention, only the tDCS group retained gains one month after training. Furthermore, a subset of studies has failed to observe any cognitive benefits from tDCS (Nilsson et al., 2017). Such discrepancies in findings may indicate that responses to tDCS could be highly influenced by aspects such as different study protocols (e.g., training dose, stimulation parameters, etc.) and interindividual variability (e.g., educational background, baseline WM capacity, etc.). Regarding the latter, some studies suggest that tDCS benefits are more pronounced in individuals with higher levels of education (Berryhill & Jones, 2012; Stephens & Berryhill, 2016) or higher baseline WM (Katz et al., 2017). Conversely, others posit that only those with lower baseline performance levels experience significant benefits from tDCS, due to their greater potential for improvement (Arciniega et al., 2018; Au et al., 2022; Krebs et al., 2021). This contradictions highlight the need for further exploration of factors mediating the effectiveness of tDCS in older adults.

Despite the variability in behavioural outcomes, neuroimaging studies provide more consistent support for the neural impact of tDCS in older populations (Antonenko et al., 2018; Cespón et al., 2017; Meinzer et al., 2013; Nissim al., 2019a; Nissim al., 2019b; Vecchio et al., 2021) However, few studies have specifically investigated tDCS-induced changes in WM-related oscillatory activity in older adults, even though understanding these neurophysiological mechanisms is essential for elucidating the effects of tDCS (Emonson et al., 2019; Habich et al., 2020). The existing literature, though limited, has yielded promising results. For instance, Zaehle et al. (2011) reported that a single tDCS session enhanced theta and alpha power during WM tasks in younger adults. Similarly, studies on resting-state brain activity in younger populations showed significant increases in alpha and beta power (Mangia et al., 2014) and in theta power (Miller et al., 2015) following tDCS. In older adults, Vecchio et al. (2021) observed decreases in delta and theta power and an increase in alpha power in resting-state activity after a 13-minute tDCS session. Such findings suggest that tDCS may modulate oscillatory dynamics relevant to WM performance. Additionally, Jones et al. (2017, 2020) reported that WM improvements following tDCS were associated with a significant reduction in posterior alpha power and an increase in the synchrony between frontal theta and posterior low alpha activity, further supporting the role of oscillatory mechanisms in tDCS efficacy.

Nevertheless, given the current knowledge gaps and inconsistent findings, further research is needed to explore the modulatory effects of tDCS across distinct phases of the WM process in older adults. Most studies to date have focused on tasks such as the N-back, which do not allow for a precise temporal distinction between different WM phases, complicating the interpretation of results (Haque et al., 2021; H. Kim, 2019; Román-López et al., 2019). For instance, in the study by Jones et al. (2017), changes in theta and alpha bands were examined only during the delay period. A more comprehensive investigation of WM phases—encoding, maintenance, and retrieval—could inform optimisation of tDCS protocols in older adults, enabling targeted interventions tailored to specific WM processes (Cespón et al., 2017; Emonson et al., 2019; Habich et al., 2020).

In the current study, we adapted two widely validated WM tasks—the Letter Span (verbal-auditory modality; Crannell & Parrish, 1957) and the Corsi Block-Tapping Test (visuo-spatial modality; Corsi, 1972)—and included distinct encoding, maintenance, and retrieval phases. Furthermore, each task included two conditions: a retention condition requiring participants to retain information and a manipulation condition requiring participants to manipulate information. This approach allowed for a detailed examination of different WM components. Additionally, we employed electroencephalography (EEG) to measure oscillatory power throughout the entire WM process. The objective of this study was to determine whether a WM training programme consisting of six sessions over three weeks, combined with tDCS, would enhance WM performance and neural activity in older adults. We hypothesised that both the experimental and control groups would benefit from the WM training, but that the experimental group receiving tDCS would show significantly greater improvements in verbal WM tasks both during training (learning effects) and in subsequent tests (near-transfer effects) compared to the control group (receiving cognitive training plus sham tDCS). As stimulation was applied to the left DLPFC, which has been shown to be more effective in enhancing verbal WM (Wischnewski et al., 2024), we expected that these improvements would correlate with increases in theta and alpha power. Furthermore, we anticipated sustained gains in the experimental group, with long-lasting effects observed 6–8 weeks post-training, particularly in the verbal domain. Lastly, we investigated the impact of individual differences (e.g., educational level, baseline WM performance, and problem-solving skills) on the efficacy of tDCS.

## METHODS

### Participants

A power analysis was conducted using G*Power 3.1 (Faul et al., 2009) to estimate the sample size required to detect significant differences between the tDCS and the control group using a Repeated Measures ANOVA, specifically for within – between interactions. The parameters were set as follows: effect size f = 0.25 (small), alpha level = 0.05, and desired statistical power = 0.8. The analysis indicated that a minimum sample size of 34 participants (17 per condition) was necessary to achieve sufficient power.

A total of 42 healthy older adults (24 women, 18 men, mean age = 68.80 years, SD = 4.91, range = 62–80 years) participated in the study. Participants were recruited from the local community through public announcements. Exclusion criteria included left-handedness, metal implants in the head or skull, electronic devices in the body (e.g., pacemakers), a history of psychiatric or neurological disorders (including epilepsy and seizures), a history of alcohol or drug abuse, vision and/or hearing impairments, previous surgical procedures involving the head or spinal cord, or taking medications that might influence cognitive abilities (e.g., those causing reduced attention, fatigue, or memory impairment).

Participants were first screened via telephone to ensure they did not meet any of the exclusion criteria. Eligible participants were invited to an introductory session where their eligibility was further confirmed using the Montreal Cognitive Assessment (MoCA), the Geriatric Depression Scale (GDS), and a medical screening questionnaire that included a detailed review of their clinical history. At this stage, participants were excluded if they scored below 26 points on the MoCA, more than five points on the GDS, or met any other exclusion criteria from the medical screening. Four participants were excluded due to failing the MoCA. During this session, eligible participants received a comprehensive description of the study, including detailed information about tDCS and EEG. After providing this information, participants signed an informed consent form. The final debriefing, including allocation to the study groups, was conducted at the end of the study. Participants were divided in two groups: tDCS and Control (sham). The groups were matched based on MoCA scores, age, sex, education level, and highest degree obtained. Before each testing session, participants confirmed that they had slept for at least six hours the previous night, refrained from alcohol or drug use within eight hours prior, and abstained from coffee consumption within two hours of testing. Five participants did not complete the training and were excluded from the final analysis. Ethical approval for this study was granted by the Ethics Research Committee of Edge Hill University.

### Experimental procedure

The study consisted of ten sessions: an initial screening session to confirm eligibility, a pre-training session, six training sessions spaced 24 to 72 hours apart, a post-training session, and a follow-up session 6–8 weeks later (see Figure 1). Participants completed at least two training sessions per week, and all sessions were completed within three weeks. The post-training session was conducted within 24 to 48 hours of the final training session. Four participants did not attend the follow-up session.

**Figure 1.**
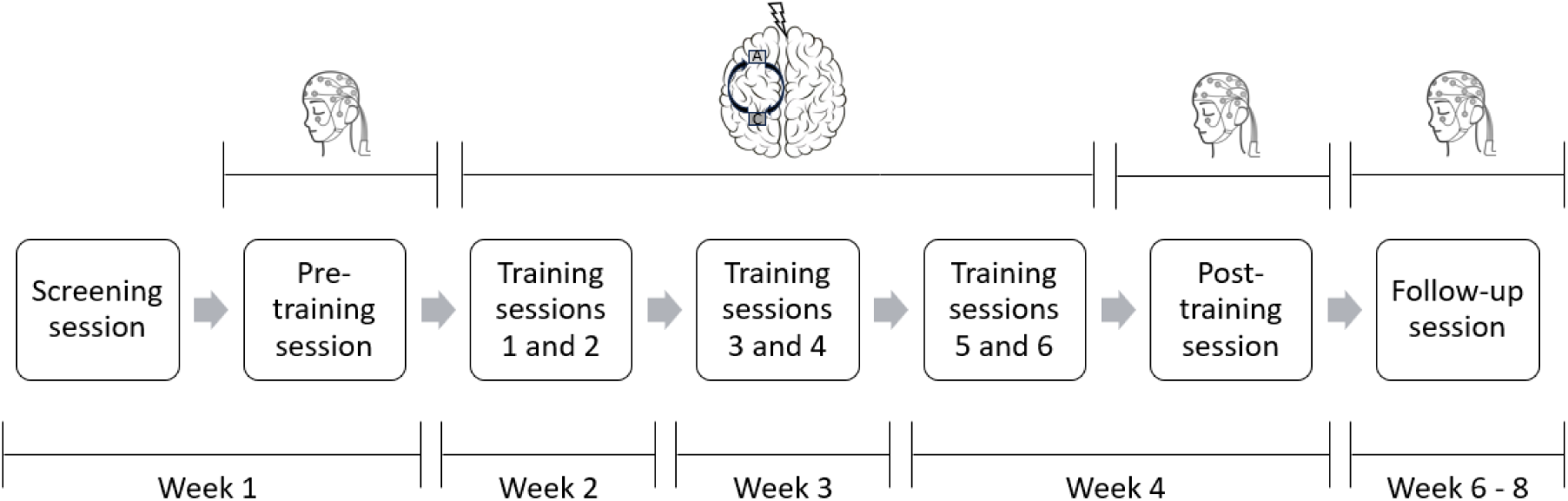
Participants completed ten sessions: one screening session, one pre-training, six training, one post-training, and one follow-up session. EEG was recorded during the pre-, post-, and follow-up sessions. tDCS was delivered during the first 20 minutes of every training session.

Before each session, participants completed a brief questionnaire assessing their current state, including factors such as sleep, medication intake, recent exercise, alcohol consumption, motivation, and fatigue. EEG data were collected during the testing sessions, while tDCS was applied for the first 20 minutes of each training session. Testing sessions lasted approximately 2.5 hours, and training sessions lasted one hour.

### Training tasks

WM training was delivered through two tasks: an adapted version of the visuospatial N-back and an adapted version of the Operation Span (OSPAN). Both tasks were presented using PsychoPy Software (Peirce, 2007) and modified to suit the study’s requirements. Participants in the experimental group received stimulation during the OSPAN task only, with no stimulation during the N-back task. This was the case because stimulation was applied to the left hemisphere, which has been shown to be more effective in enhancing verbal WM (Wischnewski et al., 2024).

### OSPAN

The OSPAN combined features of the original test (Turner & Engle, 1989) with a backwards version of the Digit Span (Figure 2A) to reduce potential learning effects and prevent the use of similar stimuli to the Letters Span. In this task, participants listened to a sequence of digits and then recalled them in reverse order while solving complex arithmetic questions. For the arithmetic questions, participants were presented with a complex operation and an answer, and they had to decide if the provided answer was correct or incorrect by clicking “Yes” or “No” on the screen with the computer mouse. They had a maximum of six seconds to respond, and feedback was given immediately, with “Correct” or “Incorrect” appearing on the screen after each answer. Each arithmetic question was followed by a digit, and after the final digit, participants typed their answer using the keyboard.

**Figure 2.**
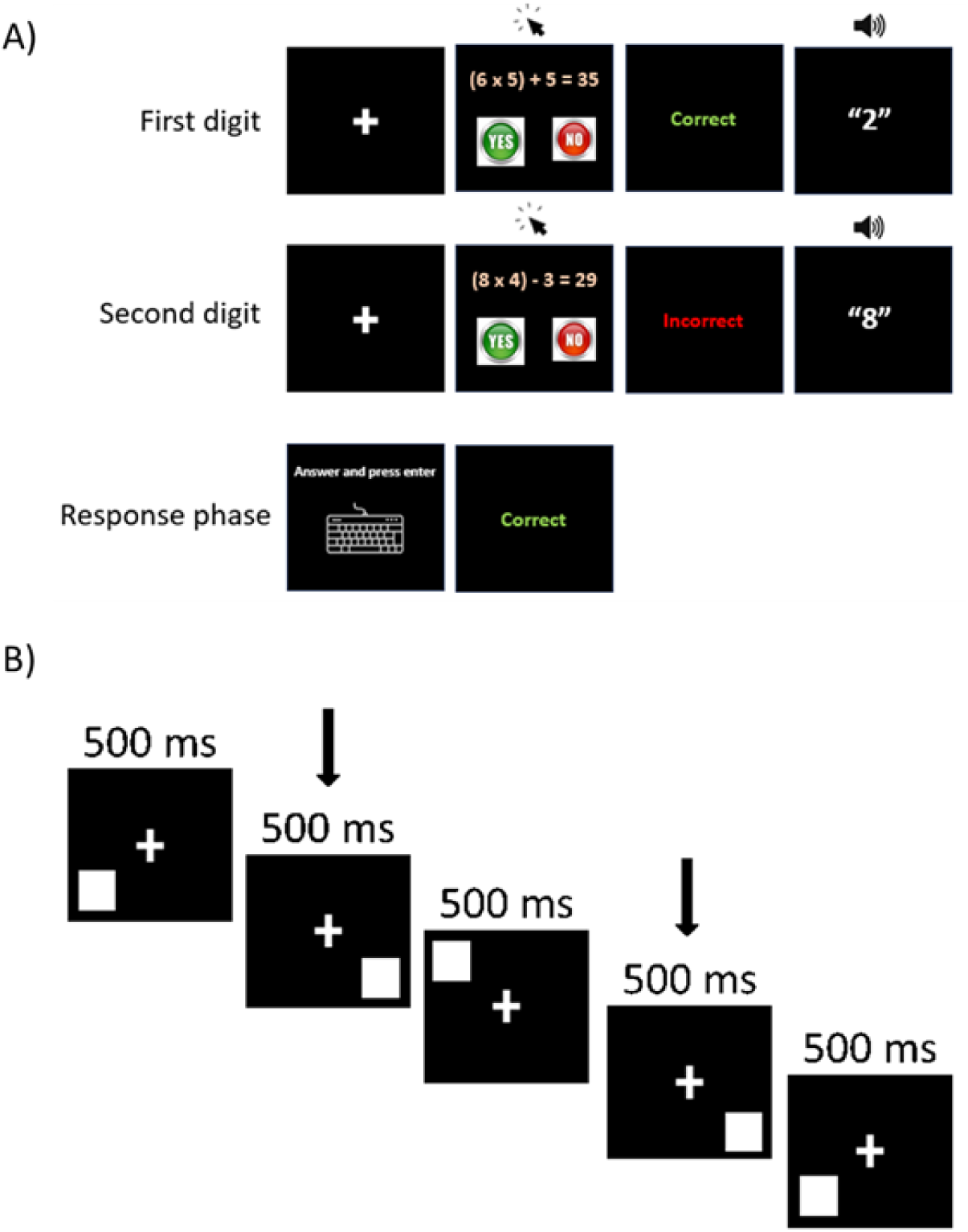
A) Example of the adapted version of the OSPAN. Participants would solve an arithmetic question followed by a digit (auditory). Their goal was to recall the digits in backwards order using the keyboard while solving the arithmetic questions correctly. B) Example of a 2-back trial. The arrows show the squares that must be compared. In this case, the participant would have to press “a” because squares appeared in the same location. In both tasks participants received feedback of their performance after every trial (OSPAN) or block (N-back). According to their performance the difficulty of the task would increase or decrease.

During the response phase, participants had no time limit but were not allowed to correct typing mistakes. This ensured that participants typed the digits in the correct order rather than randomly to avoid forgetting them before typing them in the correct order. At the end of each trial, feedback indicated whether the digits were recalled correctly with the words “Correct” or “Incorrect”. All participants practised the tasks separately and then together before beginning the training. This practice was repeated before each training session as a “warm-up”. The warm-up included two trials of two digits without maths problems, followed by two trials of only arithmetic questions without a time limit, and finally four trials of the combined tasks. Training commenced only after participants were familiar with the task.

Each training session consisted of three blocks of ten trials each, and all participants started with a series of two digits. The number of digits increased or decreased based on performance. If a participant recalled the digits correctly twice in a row at the same level, another digit was added to the series for at least the next two trials. Conversely, if they answered incorrectly twice in a row at the same level, a digit was removed from the series for at least the next two trials. If they had mixed results (one correct and one incorrect, or vice versa), they remained at the same level until one of the other scenarios occurred.

Two metrics were extracted for each session: raw scores and the highest level completed (Conway et al., 2005). Raw scores were calculated by multiplying the number of correct answers by their respective level and then adding up the scores of all the blocks. The highest level completed was defined as the highest level where participants correctly recalled the two series of digits. The task took approximately 20 minutes to complete.

### N-back

The N-back (see version of Au et al., 2016) involved displaying a series of white squares on a black screen in eight different possible positions (Figure 2B). The squares appeared one by one for 500ms, and participants had to indicate whether the last square was in the same location as the square that appeared “n” trials before. If the square was in the same location, participants pressed “a”; if it was not, they pressed “l”. Participants were instructed to respond every time a square appeared and to rely on their memory. They had 2000ms to answer before the next square appeared.

The training consisted of 14 blocks of 20 trials each. Seventy percent of the trials (14) were incongruent (no match in locations), and thirty percent (6) were congruent (match in locations). Before starting the training, participants completed four short practice blocks of 10 trials each to familiarise themselves with the task. The first two blocks were 1-back, and the last two were 2-back. After each block, participants received feedback on their score, which reflected the percentage of correct answers (e.g., if a participant answered 10 trials correctly, the score would be 50%). Based on their score, the difficulty of the following block was adjusted. If a participant scored 90% or higher, the level increased by one (e.g., from 1-back to 2-back). If they scored between 71% and 89%, they remained at the same level. If the score was 70% or lower, they dropped one level (e.g., from 3-back to 2-back), except for 1-back, where they stayed at the same level. These percentages ensured that participants relied on their memory skills and did not advance by chance.

Participants began each training session with 1-back. Two metrics were extracted for each training session: the raw score and the highest level reached (e.g., 2n, 3n, 4n, etc.). Scores were calculated by multiplying the number of correct answers by the level of the respective block and summing the scores of all the blocks. The task took approximately 20 minutes to complete.

### Working Memory tests

#### Letter Span

Two WM tests were employed, both presented using PsychoPy Software (Peirce, 2007). One of the tests was an adapted version of the Letter Span (Figure 3A), used in previous studies (e.g., (Pavlov & Kotchoubey, 2017; Postle et al., 2006; Scheeringa et al., 2009), but in the auditory modality. The test commenced with a fixation cross displayed for one second, followed by a sequence of six letters heard one by one, each separated by approximately one second. The test included twelve consonants to avoid forming words: C, F, H, J, L, N, K, P, Q, R, V, W.

**Figure 3.**
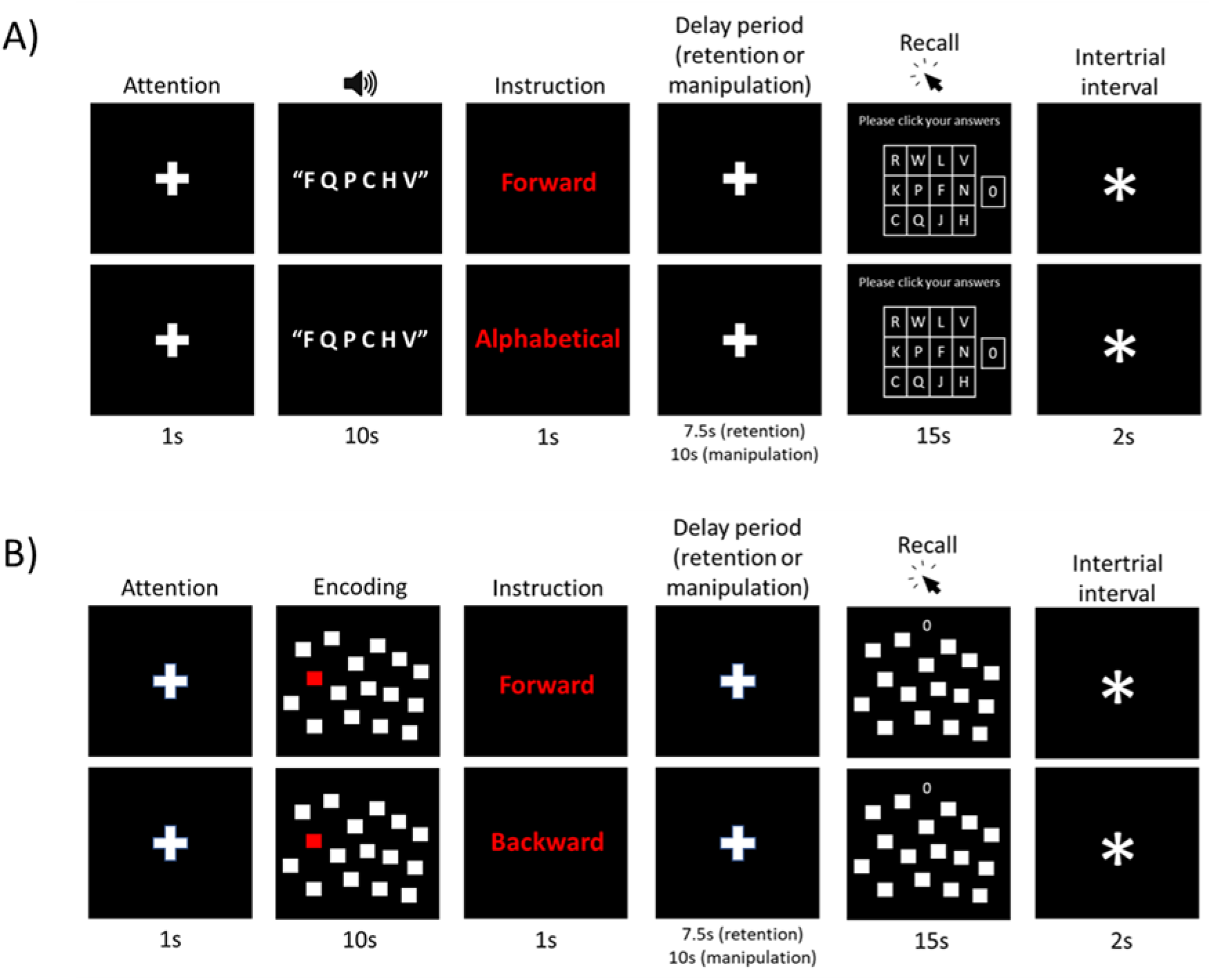
A) Adapted version of the Letter Span. A sequence of six letters was heard one by one followed by an instruction with the order in which they had to be recalled. Then, they would have a delay period followed by a grid to click the relevant letters in the correct order. B) Adapted version of the Corsi Block-Tapping test. A sequence of six squares were coloured one by one followed by an instruction with the order in which they had to be recalled. Then, they would have a delay period followed by a group of white squares displayed on the screen to click the relevant ones in the correct order.

After participants heard the last letter, an instruction appeared showing either “Forward” (retention condition) or “Alphabetical” (manipulation condition) in red font, followed by another fixation cross to mark the beginning of the delay period. This delay lasted seven and a half seconds for the “Forward” instruction (retention condition) and ten seconds for the “Alphabetical” instruction (manipulation condition). In the retention condition, participants had to remember the sequence of letters in the exact same order in which they were displayed, whereas in the manipulation condition they had to reorder the letters according to the alphabet in ascending order. After the delay period, to indicate the start of the recall phase, a 4 x 3 grid displaying the potential letters appeared on the screen with an additional cell containing a zero. Participants were instructed to click the letters with the mouse as quickly as possible in the specified order according to the instruction. They were allowed to use the zero to fill in a space in the sequence if they forgot a letter, but only once per trial. The letters always appeared in the same positions on the grid to avoid requiring visual searching skills. This recall phase lasted 15 seconds, followed by a 2-second intertrial interval before the next trial began.

To familiarise themselves with the task, the use of the mouse, and the letter locations on the grid, all participants completed one practice block consisting of three trials of each condition. If the researcher observed that the instructions were still unclear, an additional practice block was provided. Excluding the practice trials, the test comprised eight blocks of six trials each (24 trials per condition). Each block included three randomised trials of each condition. Participants earned one point for each correctly recalled letter in the sequence, with a maximum score of six points per trial. This scoring method was chosen to keep participants focused and motivated, even if they missed some letters in a sequence.

#### Corsi Block-Tapping test

The second test was an adapted version of the Corsi Block-Tapping test (Figure 3B) that followed a similar design as the adapted version of the Letter Span. In this case, instead of letters, sixteen white squares were appearing on the screen and six of them were coloured in red, one by one, for approximately one second each. After the last square was coloured, an instruction appeared showing either “Forward” (retention condition) or “Backward” (manipulation condition) in red font, followed by a fixation cross to mark the beginning of the delay period. This delay lasted seven and a half seconds for the “Forward” instruction and ten seconds for the “Backward” instruction. In the retention condition, participants had to remember the sequence of squares in the exact same order in which they were displayed, whereas in the manipulation condition they had to reorder the squares backwards. After the delay period, the same white squares that were shown during the encoding phase appeared again on the screen with a zero on the upper central part. Participants were instructed to click the squares with the mouse as quickly as possible in the specified order according to the instruction. They were allowed to use the zero to fill in a space in the sequence if they forgot a square, but only once per trial. This recall phase lasted 15 seconds, followed by a 2-second intertrial interval before the next trial began.

To familiarise themselves with the task, the use of the mouse, and the squares’ locations on the grid, all participants completed one practice block consisting of three trials for each condition. If the researcher observed that the instructions were still unclear, an additional practice block was provided. Excluding the practice trials, the test comprised eight blocks of six trials each (24 trials per condition). Each block included three randomised trials of each condition. Participants earned one point for each correctly recalled square in the sequence, with a maximum score of six points per trial.

The order of the tests was counterbalanced across participants, and the tests took approximately 30 minutes each to be completed. The total duration of a session (including the EEG set-up) was between 2h and 2.5hrs. Participants had the opportunity to take breaks both between tests and within them (between blocks). The duration of the breaks was variable and depended on the participant. Before recruitment, both tests were piloted with a group of ten old adults and five young adults to adjust for difficulty factors such as the number of elements to recall, the duration of the phases, and the answer method. Moreover, stimuli were also tested. For instance, participants had difficulties differentiating the “P” from the “T” or the “F” from the “S” thus the “T” and the “S” were excluded.

During the EEG set-up, older participants completed the short version of the Everyday Problem-Solving Test (EPT; Willis, 1996) which included 28 open questions related to daily living scenarios (two questions per scenario) where information (e.g., labels or charts) to answer the questions was provided. This test has been used previously in studies with older adults like the ACTIVE study (Tennstedt & Unverzagt, 2013) and it has been proved to predict real functioning in individuals as its outcomes have been compared with means of direct behavioural observation (Moore et al., 2007; Schmitter-Edgecombe et al., 2011). Although the test has been primarily validated with US American samples, we decided to include all the questions as the information provided was sufficient for most of the participants to answer the questions correctly and, when we excluded the questions more culturally related to the USA, results did not change significantly. The time of completion of test was 15 - 20 minutes approx.

### tDCS protocol

Stimulation was delivered through a battery driven HDCstim (MagStim Company Limited; Whitland, U.K.). In the tDCS condition, an amplitude of 2mA was applied during the first 20 minutes of every training session with a 30-seconds linear ramp up at the beginning and a 30-seconds linear ramp down at the end to reduce discomfort. In the sham condition, stimulation began with 30-seconds linear ramp up followed by 30 seconds of stimulation at 2mA and then a 30-seconds linear ramp down. This method was used to create sensations like those of the real stimulation (Ambrus et al., 2012; Russo et al., 2013). 5 x 5cm conductive rubber electrodes inside sponge soaked in a 0.9% isotonic saline solution and conductive gel to increase conductivity were placed on the scalp. The anodal electrode was placed over the left DLPFC and the cathodal electrode was placed over the left Parietal Cortex (anode on F3 – cathode on P3 according to the international 10 – 10 system). Electrodes were held in place using a 64-channel cap (Brain Products GmbH) that was also used to identify the target locations. Impedance was kept under 5 kΩ.

### EEG data acquisition

EEG data was acquired using a 64-channel actiCAP active electrode array and an actiCHamp amplifier system (Brain Products GmbH) at a 1000Hz sampling rate. Electrodes were placed over the scalp following the international 10 – 10 system. EEG data were online referenced to Cz electrode. Electrodes’ impedances were kept below 10 kΩ to minimise noise and ensure a better quality of the EEG signal. BrainVision Recorder Software was used to record the signal. Data was collected continuously throughout the execution of the tests.

### EEG data pre-processing

EEG data were pre-processed to reduce the signal-to-noise ratio using the EEGLAB toolbox (Delorme & Makeig, 2004) in MATLAB (R2023b, The Mathworks, USA). Data were resampled at 512Hz and bandpass filtered at 1Hz – 40Hz. An artefact with a narrow peak at 20Hz likely due to interference caused by an external electronic device was removed using the *cleanline* EEGLAB plugin. Visual inspection was then conduced to remove bad segments of data which involved mainly cutting long breaks. Bad channels rejection was performed using the function *pop_clean_rawdata*. Channels were rejected if they met any of the following criteria: 1) they were flat for five seconds or more, 2) they were not correlated to neighbouring channels by at least 80%, or 3) their amplitudes were +/- 4 SD out of range. Although this would be normally the rule, the final decision of what channels to reject was made by the researchers according to their criteria (not rejecting more than 10% of the channels). The next step was to correct for blinking and muscular artifacts using independent component analysis (ICA). Components were rejected, with the help of the function *pop_iclabel*, if they met any of the following criteria: 1) they contained more than 80% of muscular activity or 2) they contained more than 80% of eye activity. Like with the rejection of channels, the final decision of which components to reject was made by the researchers according to their criteria (not rejecting more than 20% of the components).

Finally, rejected channels were interpolated, and the data were re-referenced to the average reference. Epochs were then extracted without baseline correction for each WM phase. Only trials with a minimum of 50% accuracy (three correct answers out of six in the sequence) were included in this analysis. Participants who had less than 25% valid trials per test condition (six out of twenty-four) were excluded from the analysis of that specific condition.

The encoding epochs were stimulus-locked (relative to stimulus onset) from 0 to 8.5 seconds. Then, after every phase, a trigger marked the beginning of a new phase with the following durations: delay-retention [0 to 7.5 seconds], delay-manipulation [0 to 10 seconds], recall-retention [0 to 4 seconds], and recall-manipulation [0 to 4 seconds]. For example, in the Letter Span, the encoding phase commenced with the first letter of the sequence one second after the initial fixation cross appeared on the screen. Following the presentation of the final letter in the sequence, a fixation cross signalled the onset of the delay period, marking the beginning of the maintenance phase. Subsequently, the recall phase was initiated by the appearance of the response grid on the screen. The duration of epochs was according to the length of the phase except for the recall phases where it was decided to use a window of four seconds to maintain the duration of the trials consistent across participants. The reasons for this decision were the high variability between participants response times and the high number of trials excluded due to short duration (less than five seconds).

After this, epochs were segmented into smaller sub-segments of 1-sec and trials that contained strong residual artifacts were rejected by eye. If after visual inspection a participant had less than 15 valid sub-segments of data in the maintenance (retention or manipulation) and/or recall phases in any of the test conditions, the participant was excluded only from the analysis of that specific condition. According to this criterion, in the Letter Span, no participants were excluded from the retention condition, but two participants from the tDCS group were excluded from the manipulation condition. Regarding the Corsi Test, three participants from tDCS group and four participants from the Control group were excluded from the retention condition, and nine participants from the tDCS group and four participants from the Control group were excluded from the manipulation condition. One participant from the Control group was excluded from all the tasks due to researcher’s errors during recording of neurophysiological data.

### Spectral analysis

Spectral analysis was conducted using the toolbox Fieldtrip (Oostenveld et al., 2011). Spectral power was extracted using a Fast Fourier Transformation (FFT) with a Hanning taper focusing on frequencies from 2Hz to 40Hz. Both WM tests and their conditions were analysed separately. Thus, there were in total four parallel analyses: retention – Letter Span, manipulation – Letter Span, retention – Corsi Test, manipulation – Corsi Test.

### Statistical Analysis

Statistical analysis of behavioural results was conducted using RStudio (2023). To explore the effects of the stimulation on the training, Linear Mixed Effects Models (LMEM) were used with condition and session as fixed factors and participants as the random effect. Two models per training task were created with raw scores and maximum level achieved per session (maximum number of elements correctly recalled) as dependent variables.

To examine the effects of the tDCS in the WM tests, the scores of the pre- and post-training sessions were compared using a LMEM including condition and session as fixed factors and participants as the random effect. Similarly, we analysed the long-lasting effects of tDCS using a LMEM including condition and session as fixed factors and participants as the random effect, but this time the scores of the post-training session and the follow-up sessions were compared.

Both for training tasks and WM tests, in cases where significant main effects or interaction were found, post-hoc analysis using Tukey correction for multiple comparisons and a Kenward-roger degrees-of-freedom method were used to explore those effects further. To confirm the fit of the models Q-Q plots, histograms of residuals and the Shapiro-Wilk normality test of residuals were used.

To explore the effects of individual differences in the training tasks, Likelihood Ratio Tests were conducted to identify whether highest degree of education would improve the original models (condition + session). Moreover, univariate ANOVAs were conducted to examine the effects of score at baseline with condition and baseline score (score in session 1) as predictors, and the result of subtracting the score of session 1 from the score of session 6 as the outcome variable. Regarding WM tests, Likelihood Ratio Tests were conducted to identify whether highest degree of education would improve the original model (condition + session). Furthermore, two separate univariate ANOVAs were conducted to identify any effects of baseline score or problem-solving skills. The first ANOVA included condition and the score of the pre-training session (score at baseline) as predictors, and the second one included condition and the score of the EPT as predictors. The outcome variable used in these two analyses was obtained by subtracting the score of the pre-training session from the score of the post-training session.

To establish a link between any potential improvements during training and the WM tests, LMEM were conducted for each WM test with session and conditions as fixed factors, participant as random factor, and improvements in raw scores and max level of the training tasks as covariates (separate models for each covariate). The values used for the improvements in raw scores and max level of the training tasks were obtained by subtracting the outcomes of the first training session from the last training session. The OSPAN was matched with the Letter Span and the N-back was matched with the Corsi Test.

The EEG data were analysed using Fieldtrip in MATLAB. Before conducting the statistical analysis, power spectrum was averaged by trials to have a 2D matrix (channel x frequency) per condition (Control or Experimental) and per participant. Data were visualised with boxplots for every WM phase and frequency band. Following Klimesch (1999), the ranges of the frequency bands were defined using the individual peak alpha frequency (IAF) from the traditional alpha band (8 – 12Hz) for every participant at every phase of the WM process. The alpha band was divided in two: low – alpha, from the IAF to two frequencies (Hz) below the IAF, and the high-alpha, the two immediate frequencies (Hz) above the IAF. The theta band was defined as five frequencies (Hz) below the IAF to one frequency below the lower bond limit of the low-alpha band (Klimesch, 1999). Power was log-transformed to improve sensitivity, robustness and reduce impact of high variance between participants.

To compare spectral power between the pre-, post- and follow-up sessions, cluster-based permutation independent sample t-tests were used in all three frequency bands. This method was chosen because it controls for the multiple comparisons problem (Maris & Oostenveld, 2007; Meyer et al., 2021). Two types of variables were used for the comparisons: absolute power at every WM phase (encoding, delay-retention, delay-manipulation, recall-retention and recall manipulation), and changes in power from one WM phase to another (reflecting relative changes). Relative change was calculated by subtracting the power of one WM phase from the other. For example, power spectrum in the encoding phase was subtracted from the power spectrum in the delay-retention phase and that value was used for the Montecarlo permutations. In total there were six comparisons: encoding vs delay-retention, delay-retention vs recall-retention, encoding vs recall-retention, encoding vs delay-manipulation, delay-manipulation vs recall-manipulation and encoding vs recall-manipulation). Analyses were conducted using the function ft_freqstatistics. The highest sum of t-values across all clusters in each permutation were taken (10,000 iterations) and added to a distribution of cluster statistics. Effects observed in the data were compared against this cluster distribution. A p value of 0.05 was considered as the threshold for significance.

To calculate effect sizes, Meyer et al. (2021) guidelines were followed which state that providing an upper and lower bound for the Cohen’s *d* is more informative. Using Fieldtrip, the upper bound of the effect size was calculated using the maximum effect within the significant cluster and the lower bound was obtained using a rectangular shape to fit the cluster and extracting the averaged data from it.

Finally, Spearman correlations were performed in Python (3.10) within a Spyder environment to identify associations between neurophysiological data and behavioural improvements between sessions. Variables were created by grouping channels with frequency bands and WM phases (e.g., Enc_Cz_Theta). Statistical analyses were conducted separately for each frequency band and for each stimulation condition. The False Discovery Rate (FDR) was used to adjust p-values and correct for multiple comparisons.

## RESULTS

### Training results

#### OSPAN

For the OSPAN, both LMEMs (raw scores and max level achieved) showed no significant interaction between condition and session. However, a significant main effect of session was observed in both models (raw scores: *F*(5, 155) = 35.09, *p* < .001, *η^2^* = 0.53; max level achieved: *F*(5, 155) = 26.08, *p* < .001, *η^2^* = 0.46). No significant main effect of condition was noted. The model analysing raw scores had an AIC = 1603.22, with the variance explained by the fixed factors at 0.15 (marginal R^2^) and the variance explained by both fixed and random effects at 0.84 (conditional R^2^). For the max level achieved, the AIC was 482.2, with the variance explained by the fixed factors at 0.11 (marginal R^2^) and the variance explained by both fixed and random effects at 0.84 (conditional R^2^). The normality test was non-significant for the raw score model (W = 0.99, *p* = 0.08), but significant for the max level model (W = 0.98, *p* = 0.03), yet the results were deemed reliable due to the large sample size and similarity in outcomes across models.

Post-hoc analysis of the results of the raw scores and max level showed that there were no significant differences in performance between groups at baseline (session 1) in either of the types of scores (raw scores: B = −80.29, SE = 49.9, *t*(49.6) = −1.61, *p* = .9; max level achieved: B = −0.3, SE = 0.29, *t*(74.9) = −1.04, *p* = 1). In raw scores, participants improved significantly across sessions (session 1 vs session 6: B = −27.89, SE = 2.49, t(155) = −11.2, *p* < .001) with the first significant increase at session 2 (session 1 vs session 2: B = −9.44, SE = 2.49, *t*(155) = −3.79, *p* < .01), a second significant increase at session 4 (session 2 vs session 4: B = −8.32, SE = 2.49, *t*(155) = −3.34, *p* = .01) and a third significant increase at session 5 (session 4 vs session 5: B = −7.49, SE = 2.49, *t*(155) = −3.19, *p* < .05) (see Figure 4A). In max level achieved, participants also improved significantly across sessions (session 1 vs session 6: B = −1.33, SE = 0.15, *t*(155) = −9.05, *p* < .001), with the first significant increase at session 3 (session 1 vs session 3: B = −0.76, SE = 0.15, *t*(155) = −5.17, *p* < .001), and the second one at session 5 (session 3 vs session 5: B = −0.57, SE = 0.15, *t*(155) = −3.9, *p* < .01) (see Figure 4B).

**Figure 4.**
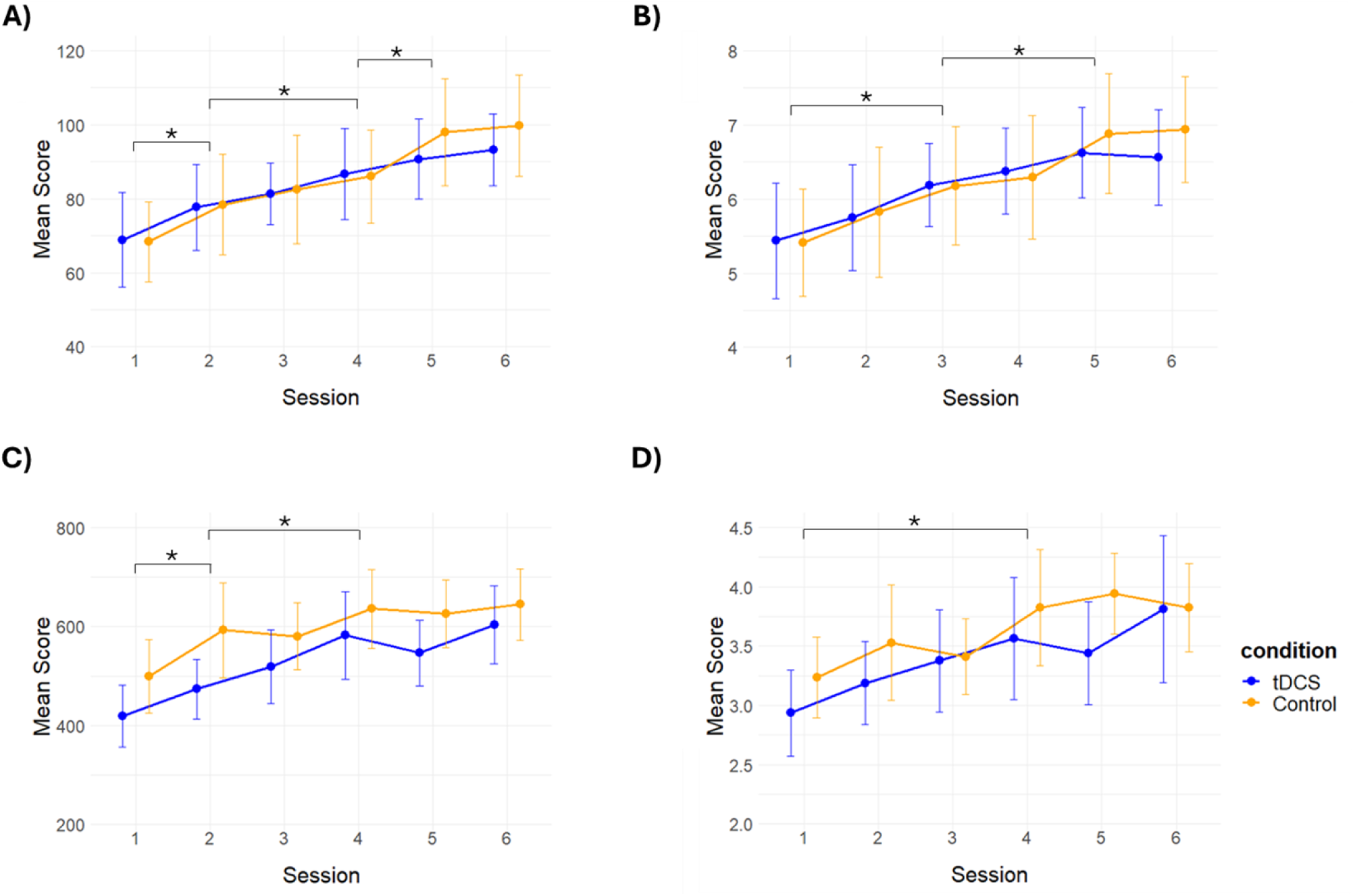
Mean scores and confidence intervals of training sessions: OSPAN and N-back. A) Raw scores – OSPAN. B) Max level achieved – OSPAN. C) Raw scores – N-back. D) Max level achieved – N-back. The * mark significant improvements between sessions. No significant differences between groups were found.

Regarding individual differences, the Likelihood Ratio Test showed that highest degree of education did not provide any added benefit to the initial model suggesting that that highest degree achieved was not a significant predictor of improvements across sessions. Conversely, the ANOVA conducted to explore the effects of baseline scores, evidenced a significant interaction between baseline scores and condition for raw scores (*F*(1, 29) = 7.15, *p* = .01, *η^2^* = 0.20). This effect was primarily driven by the tDCS group (*F*(1, 14) = 10.4, *p* < .01, *η^2^* = 0.43). Post hoc analysis revealed that participants in the tDCS group with lower baseline scores showed greater improvements across sessions (raw scores: B = −0.44, SE = 0.14, *t*(14) = −3.22, *p* < .01; max level achieved: B = −0.37, SE = 0.14, *t*(14) = −2.57, *p* < .05).

#### N-back

Similarly, for the N-back, both LMEMs showed no significant interaction between condition and session. However, a significant main effect of session was observed (raw scores: *F*(5, 155) = 22.66, *p* < .001, *η2* = 0.42; max level achieved: *F*(5, 155) = 8.13, *p* < .001, *η2* = 0.21). No significant main effect of condition was noted. In the model with the raw scores, the model’s relative fit was AIC = 2370.89, the variance explained by the fixed factors was 0.18 (marginal R^2^) and the variance explained by both the fixed factor and the random effects was 0.79 (conditional R^2^). Regarding max level achieved, the model’s relative fit was AIC = 411.94, the variance explained by the fixed factors was 0.11 (marginal R^2^) and the variance explained by both the fixed factor and the random effects was 0.60 (conditional R^2^). In both models the normality test was not significant (raw scores: W = 0.99, *p* = 0.3; max level: W = 0.98, *p* = 0.37).

Post-hoc analysis of the results of the raw scores and max level showed that there were no significant differences in performance between groups at baseline (session 1) in either of the types of scores (raw scores: B = −80.29, SE = 49.9, *t*(49.6) = −1.61, *p* = .9; max level achieved: B = −0.3, SE = 0.29, *t*(74.9) = −1.04, *p* = 1). In raw scores, participants improved significantly across sessions (session 1 vs session 6: B = −164.6, SE = 17.9, *t*(155) = −9.19, *p* < .001) with the first significant increase at session 2 (session 1 vs session 2: B = −73.9, SE = 17.9, *t*(155) = −4.13, *p* < .001), and second significant increase at session 4 (session 2 vs session 4: B = −76, SE = 17.9, *t*(155) = −4.24, *p* < .001) (see Figure 4C). In max level achieved, participants also improved significantly across sessions (session 1 vs session 6: B = −0.73, SE = 0.14, *t*(155) = −5.38, *p* < .001), with the only significant increase observed at session 4 (session 1 vs session 4: B = −0.61, SE = 0.14, *t*(155) = −4.46, *p* < .001) (see Figure 4D).

Individual differences in highest degree of education and baseline performance did not significantly impact results.

### Behavioural results

General Linear Models were conducted to explore differences between groups at baseline. There were no differences between conditions in the scores of the MoCA (*F*(1, 31) = 0.17, *p* = .68, *η2* = 0.006), age (*F*(1, 31) = 0.03, *p* = .86, *η2* = 0.001), sex (*F*(1, 31) = 0.33, *p* = .57, *η2* = 0.01), levels of education (*F*(1, 31) = 0.78, *p* = .38, *η2* = 0.03) and highest degree achieved (*F*(1, 31) = 1.08, *p* = .31, *η2* = 0.03). Moreover, there were no differences in WM performance at baseline between groups in any of the tests (Letters retention: *F*(1, 31) = 0.91, *p* = .35, *η2* = 0.03; Letters manipulation: *F*(1, 31) = 0.24, *p* = .63, *η2* = 0.01; Corsi retention: *F*(1, 31) = 2.44, *p* = .13, *η2* = 0.07; Corsi manipulation: *F*(1, 31) = 0.83, *p* = .37, *η2* = 0.03).

#### Letter Span – Retention

In the retention condition of the Letter Span no significant interaction between condition and session was found. Moreover, no significant main effects of session or condition were observed (see Figure 5A). The model’s relative fit was AIC = 145.38, the variance explained by the fixed factors was 0.05 (marginal R^2^) and the variance explained by both the fixed factor and the random effects was 0.84 (conditional R^2^). The normality test was not significant (W = 0.99, *p* = 0.96).

**Figure 5.**
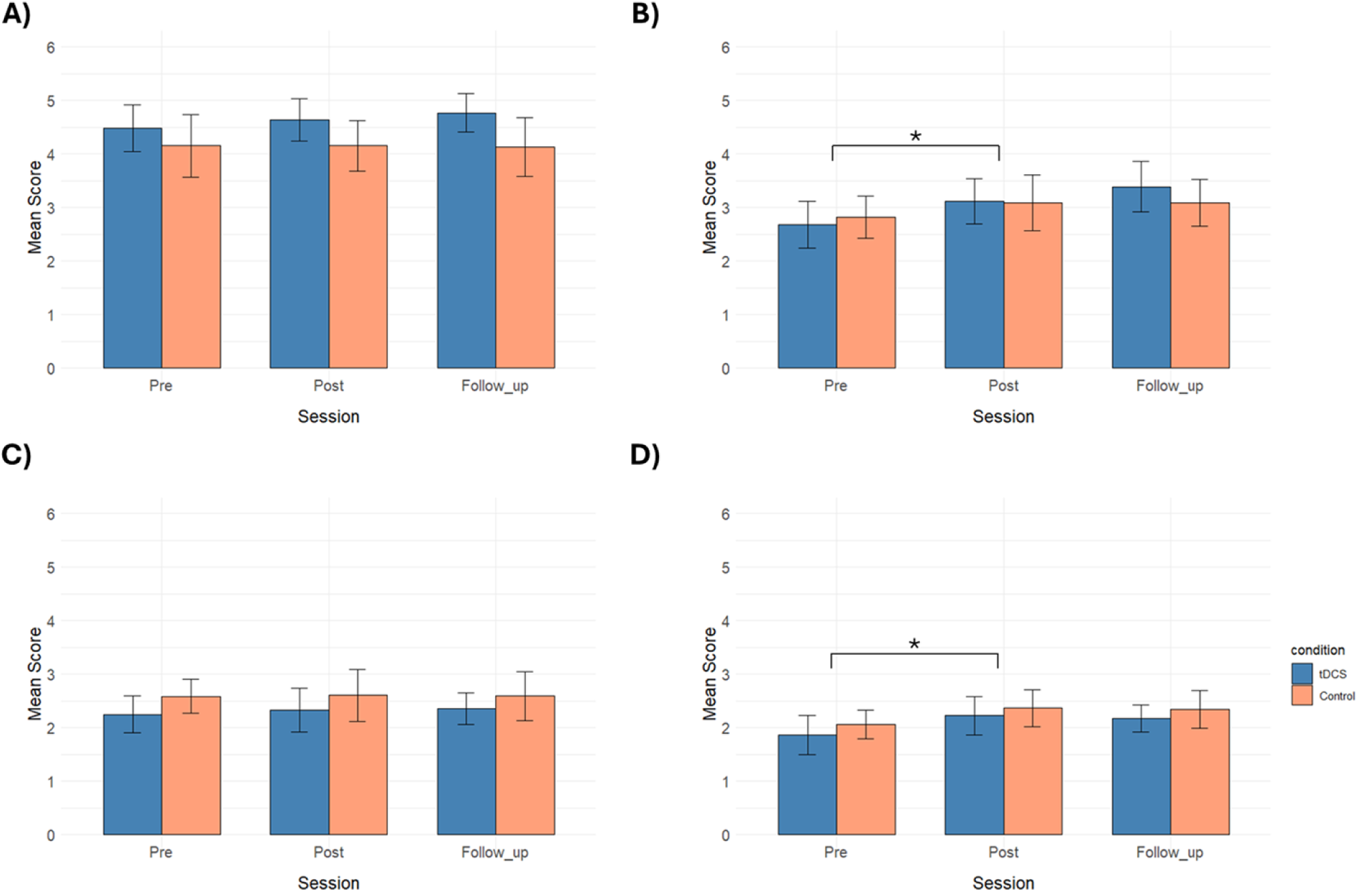
Mean scores and confidence intervals of the retention and manipulation conditions of the Letter Span and the Corsi Test. A) Retention – Letter Span. B) Manipulation – Letter Span. C) Retention – Corsi Test. D) Manipulation – Corsi Test. No significant improvements were found from pre- to post-training session in the retention condition of both tests in either of the groups. However, both groups showed significant improvements from pre- to post-training session in the manipulation condition of both tests. No significant differences were found from post-training to follow-up session in any of the test conditions (retention and manipulation) in either of the groups. These results may indicate near-transfer and long-lasting effects in the manipulation condition of both test conditions in both groups resulting from the training.

Regarding individual differences, the result of the Likelihood Ratio Test showed that highest degree of education did not improve the model significantly. Noteworthy, the variance explained by the fixed factors when highest degree was included improved (marginal R^2^ = 0.23), but the total variance explained by the model remained almost the same (conditional R^2^ = 0.85). Furthermore, ANOVA results showed no significant effect of interaction between score at baseline and condition, but a significant main effect of score at baseline was observed (*F*(1, 29) = 10.46, *p* < .01, *η2* = 0.27). Specifically, participants who performed lower at baseline showed a greater improvement from pre- to post-training session (B = −0.28, SE = 0.15, *t*(29) = −1.95, *p* = .06). Additionally, the ANOVA evidenced no significant effects of EPT.

Finally, regarding long-lasting effects, no significant effect of interaction between condition and session was found. Moreover, no significant main effect of session was found either. In the model, the variance explained by the fixed factors was 0.09 (marginal R^2^) and the variance explained by both the fixed factor and the random effects was 0.78 (conditional R^2^).

#### Letter Span - Manipulation

In the manipulation condition of the Letter Span no significant interaction between condition and session was found, however, a significant main effect of session was identified (*F*(1, 31) = 10.1, *p* < .01, *η2* = 0.25). No significant main effect of condition was observed (see Figure 5B). The model’s relative fit was AIC = 150.13, the variance explained by the fixed factors was 0.04 (marginal R^2^) and the variance explained by both the fixed factor and the random effects was 0.73 (conditional R^2^). The normality test of residuals was not significant (W = 0.98, *p* = 0.56). Post-hoc analysis showed that participants performed significantly better in the post-training session compared to the pre-training session evidencing a medium effect size and suggesting near-transfer effects resulting from the training (B = −0.35, SE = 0.11, *t*(31) = −3.17, *p* < .01, *d* = 0.42).

Regarding individual differences, the result of the Likelihood Ratio Test showed that highest degree of education did not improve the model significantly. Noteworthy, the variance explained by the fixed factors when highest degree was included improved (marginal R^2^ = 0.28), but the total variance explained by the model remained the same (conditional R^2^ = 0.73). Furthermore, ANOVA results showed no significant effects of score at baseline or EPT.

Finally, in regard to long-lasting effects, no significant interaction between condition and session was found. Moreover, no significant effect of session was observed either. In the model, the variance explained by the fixed factors was 0.01 (marginal R^2^) and the variance explained by both the fixed factor and the random effects was 0.88 (conditional R^2^). These results suggest maintenance of gains and long-lasting effects resulting from the training.

#### Corsi Test - Retention

In the retention condition of the Letter Span no significant interaction between condition and session was found. Moreover, no significant main effects of session or condition were observed (see Figure 5C). The model’s relative fit was AIC = 125.22, the variance explained by the fixed factors was 0.04 (marginal R^2^) and the variance explained by both the fixed factor and the random effects was 0.8 (conditional R^2^). The normality test of residuals was not significant (W = 0.99, *p* = 0.62).

Regarding individual differences, the result of the Likelihood Ratio Test showed that highest degree of education did not improve the model significantly. Noteworthy, the variance explained by the fixed factors when highest degree was included improved (marginal R^2^ = 0.17), but the total variance explained by the model remained almost the same (conditional R^2^ = 0.81). Furthermore, ANOVA results showed no significant effects of score at baseline. Similarly, the ANOVA evidenced no significant interaction between the EPT and condition. Nevertheless, a significant main effect of EPT was found (*F*(1, 29) = 5.78, *p* < .05, *η2* = 0.17). When exploring further the effect of EPT, results evidenced that it not a significant predictor of scores.

Finally, in regard to long-lasting effects, no significant effect of interaction between condition and session was found. Moreover, no significant main effect of session was found either. In the model, the variance explained by the fixed factors was 0.04 (marginal R^2^) and the variance explained by both the fixed factor and the random effects was 0.78 (conditional R^2^).

#### Corsi Test - Manipulation

In the manipulation condition of the Corsi Test no effect of interaction between condition and session was found, however, a significant main effect of session was identified (*F*(1, 31) = 25.81, *p* < .001, *η2* = 0.45). No significant main effect of condition was observed (see Figure 5D). The model’s relative fit was AIC = 98.01, the variance explained by the fixed factors was 0.08 (marginal R^2^) and the variance explained by both the fixed factor and the random effects was 0.84 (conditional R^2^). The normality test of residuals was not significant (W = 0.98, *p* = 0.21). Post-hoc analysis showed that participants performed significantly better in the post-training session compared to the pre-training session evidencing a medium effect size and suggesting near-transfer effects resulting from the training (B = −0.33, SE = 0.07, *t*(31) = −5.08, *p* < .001, *d* = 0.53).

Regarding individual differences, the result of the Likelihood Ratio Test showed that highest degree did not improve the model significantly. Noteworthy, the variance explained by the fixed factors when highest degree was included improved (marginal R^2^ = 0.16), but the total variance explained by the model remained almost the same (conditional R^2^ = 0.86). Furthermore, ANOVA results showed no significant effects of baseline score, but it evidenced a significant effect of interaction between the EPT and condition (*F*(1, 29) = 7.16, *p* = .01, *η2* = 0.2). That effect was mainly driven by the tDCS group (*F*(1, 14) = 4.89, *p* < .05, *η2* = 0.26) as no significant effect was observed in the Control group. No significant main effect of EPT was found.

Finally, in regard to long-lasting effects, no significant effect of interaction between condition and session was found. Moreover, no significant main effect of session was found either. In the model, the variance explained by the fixed factors was 0.02 (marginal R^2^) and the variance explained by both the fixed factor and the random effects was 0.67 (conditional R^2^). These results suggest maintenance of gains and long-lasting effects resulting from the training.

### Relationship between training and tests

In the retention condition of the Letter Span, a significant effect of the interaction between condition and improvements in raw scores of the OSPAN was found (*F*(1, 29) = 8.14, *p* < .01, *η2* = 0.22), but no significant effects of interactions involving session were observed. Specifically in the control group, participants who improved more in the OSPAN had higher scores in the retention condition of the Letter Span (*F*(1, 15) = 4.83, *p* < .05, *η2* = 0.24). This effect was not seen in the tDCS group. Moreover, no significant effects were found in any of the interactions involving improvements in max level.

Similarly, in the manipulation condition of the Letter Span, a significant effect of the interaction between condition and improvements in raw scores of the OSPAN was found (*F*(1, 29) = 9.41, *p* < .01, *η2* = 0.25), but no significant effects of interactions involving sessions were observed. Specifically in the control group, participants who improved more in the OSPAN had higher scores in the manipulation condition of the Letter Span (*F*(1, 15) = 5.75, *p* < .05, *η2* = 0.28). This effect was not seen in the tDCS group. Moreover, no significant effects were found in any of the interactions involving improvements in max level.

Regarding the Corsi Test, in the retention condition, no significant effects were found in any of the interactions involving improvements in raw scores of the N-back. Nevertheless, a significant interaction between improvements in max level and session was found (*F*(1, 29) = 4.49, *p* < .05, *η2* = 0.13). No other significant effects were found. When exploring this effect of interaction further, no significant effects were found.

Finally, in the manipulation condition of the Corsi Test, no significant effects of interactions were found in either improvements in raw scores or max level during the N-back in this task.

### Neurophysiological results

No significant changes in absolute power were observed between the pre- and post-training sessions in either group. However, significant differences in power changes across WM phases between the pre- and post-training sessions were identified.

#### Letter Span – Retention

During the retention condition of the Letter Span, the control group exhibited a significantly greater increase in theta power from encoding to the delay period in the post-training session compared to the pre-training session, specifically in the mid-frontal, central, occipital, and right parietal and parieto-occipital regions (*p* < .01, max *d* = 1.31, min *d* = 1.05). Similarly, they also demonstrated a significantly greater increase in high alpha power from encoding to the delay period in the post-training session compared to the pre-training session, specifically in the central and left parietal and parieto-occipital regions (*p* < .05, max *d* = 0.92, min *d* = 0.75). No significant differences between sessions were observed in the tDCS group.

#### Letter Span – Manipulation

During the manipulation condition of the Letter Span, the control group exhibited a marginally significant increase in low alpha power during the post-training session compared to the pre-training session, specifically from the encoding to the delay period in the left frontal, fronto-temporal, central, and parietal regions (*p* = .05, max *d* = 1.01, min *d* = 0.81). Similarly, the tDCS group also showed a marginally significant increase in low alpha power during the post-training session compared to the pre-training session, from encoding to the delay period, particularly in the occipital, right-central parietal, and parieto-occipital regions (*p* = .05, max *d* = 1.05, min *d* = 0.81). Additionally, within the tDCS group, an increase in high alpha power was observed from the delay period to recall during the post-training session compared to the pre-training session, specifically in the mid-frontal, central, and left-central parietal regions (*p* < .05, max *d* = 1.13, min *d* = 0.96).

#### Corsi Test - Retention

No significant changes in power between WM phases were observed during the retention condition from the pre- to the post-training session.

#### Corsi Test – Manipulation

During the manipulation condition of the Corsi Test, the control group demonstrated a significant decline in theta power during the post-training session compared to the pre-training session, specifically from the delay period to recall in the left frontal, central, and fronto-temporal regions (*p* < .05, max *d* = −1.03, min *d* = −0.86). Importantly, when comparing power changes between groups from pre- to post-training session, no significant differences were found in any of the frequency bands. Thus, tDCS seems to have had no impact on oscillatory activity.

#### Power changes between post-training session and follow-up session

No significant changes in absolute power or power changes across WM phases were observed between the post-training and follow-up session in either group. Moreover, when comparing power changes between groups from post-training to follow-up session, no significant differences were found in any of the frequency bands.

**Figure 6.**
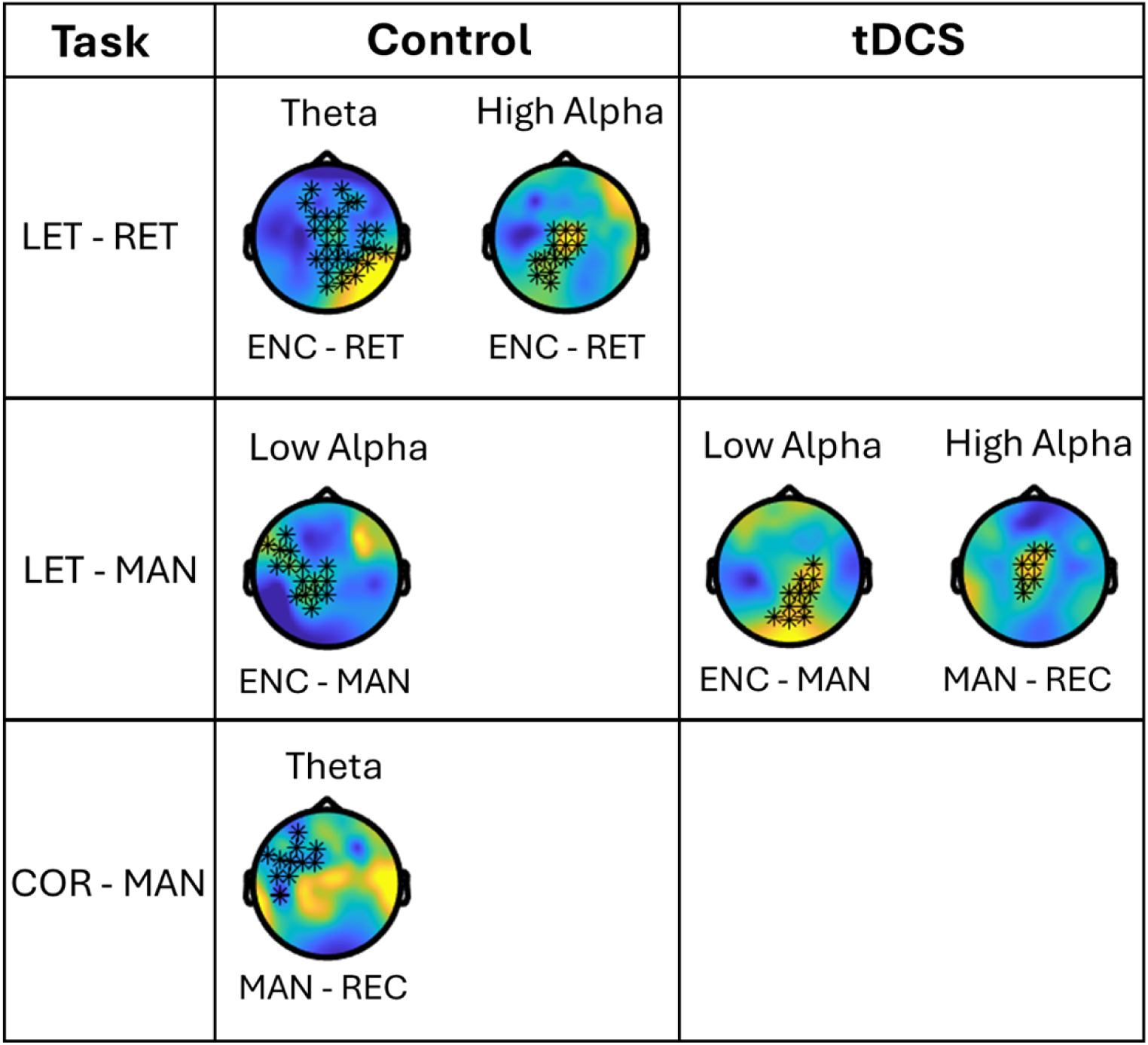
Significant differences in oscillatory power between pre- and post-training sessions. No effects were found in the retention condition of the Corsi Test.

### Correlations between EEG data and behavioural results

For the correlation analysis, each test only included those channels that were significant during the analysis of oscillatory power changes between sessions. Results showed no significant associations between changes in power between WM phases and changes in performance after correction with the FDR.

## DISCUSSION

The primary objective of this study was to investigate whether a combination of WM training and tDCS could enhance WM performance and neural functioning in older adults. We hypothesised that while both the experimental and control groups would benefit from the training, the experimental group receiving tDCS would exhibit significantly larger improvements in WM performance, evidenced by enhanced near-transfer effects, training effects, sustained long-term benefits, and more substantial gains in neural functioning.

### Training Effects

Our findings revealed that both groups demonstrated significant improvements across training sessions in the WM tasks—training effects—as hypothesised. However, contrary to expectations, no significant differences emerged between the tDCS and control groups in terms of raw scores or the maximum number of elements recalled. Similarly, in the N-back, both groups improved significantly across sessions, but no performance differences were observed between the groups. These results suggest that while the WM training was effective, the addition of tDCS did not enhance the training benefits.

This lack of tDCS-specific effects aligns with recent studies with larger samples questioning the efficacy of tDCS as an intervention for WM enhancement in older populations (Nilsson et al., 2017; Stephens & Berryhill, 2016). Nilsson et al. (2017) attributed their lack of significant findings to the incomplete understanding of optimal tDCS parameters for older populations, such as appropriate intensity and duration (Fertonani & Miniussi, 2017). They further suggested that interindividual differences, which tend to be more pronounced among ageing adults, make it particularly challenging to interpret the potential benefits of tDCS in comparison to younger populations (Heise et al., 2014; J.-H. Kim et al., 2014; Wiethoff et al., 2014). Additionally, Stephens and Berryhill (2016) suggested that longer training protocols (e.g., 10 sessions) may improve the likelihood of observing training effects. In our study, it is possible that all those factors influenced our outcomes to some extent. Specifically, we applied the same intensity and duration as Nilsson et al. (2017), identified a significant contribution of random factors in our statistical models, and employed only six training sessions, similar to the five sessions used by Stephens and Berryhill (2016).

Importantly, consistent with previous research, baseline performance effects were noticed, with participants scoring lower at baseline showing greater improvements from training (Borella et al., 2017; Bruno et al., 2024; Traut et al., 2021; Zinke et al., 2014). However, this baseline effect did not impact tDCS-related gains, further emphasising that the improvements observed were likely due to the WM training itself rather than tDCS.

### Near-Transfer Effects

Regarding near-transfer effects, no significant improvements were observed in the retention conditions of the Letter Span and Corsi Test from pre- to post-training. In the retention condition of the Letter Span this lack of improvement may be partially attributed to ceiling effects as participants exhibited high scores during the pre-training session, leaving limited room for further gains. Ceiling effects are a recognised challenge in cognitive interventions, as noted in previous research (Arciniega et al., 2018; Ball et al., 2002; Kwok et al., 2011; D. C. Park & Bischof, 2013; Shaw & Hosseini, 2020). Participants who initially performed lower did show greater improvements, which is consistent with our training results and with prior findings evidencing that baseline performance modulates the magnitude of training-related gains (Borella et al., 2017; Bruno et al., 2024; Traut et al., 2021; Zinke et al., 2014). These ceiling effects may have also masked potential tDCS benefits as demonstrated in previous studies (Krebs et al., 2021; Pergher et al., 2022; Vergallito et al., 2022). Conversely, the absence of near-transfer effects in the retention condition of the Corsi Test could partly be attributed to the training’s focus on enhancing manipulation skills, which may have facilitated backward recall but did not lead to improvements in the retention of elements per se (Blacker et al., 2017; Holmes et al., 2019; Linares et al., 2019; Morrison & Chein, 2011).

Noteworthy, in the manipulation conditions of both tests, significant improvements were observed, indicating successful near-transfer effects. These effects persisted for up to 4– 6 weeks post-training, highlighting the long-lasting benefits of the WM training intervention. However, no significant differences between the stimulation groups were detected, suggesting that tDCS did not contribute to these observed outcomes. These findings mirror those in the training tasks and align with meta-analyses reporting minimal or negligible effects of tDCS on near-transfer measures (Au et al., 2015; Karbach & Verhaeghen, 2014; Melby-Lervåg & Hulme, 2016; Summers et al., 2016). Nevertheless, the presence of near-transfer and long-lasting effects aligns with findings from previous studies and supports the claim that the training effectively enhanced specific aspects of WM (Borella et al., 2010; Brehmer et al., 2012; Holmes et al., 2019; Sala et al., 2019; Schwaighofer et al., 2015; Teixeira-Santos et al., 2019; Zinke et al., 2014).

### Correlational Findings

Correlational analysis provided additional insights but also revealed some inconsistencies. While participants who improved the most during training generally achieved higher post-training test scores, this relationship was evident only in the manipulation condition of the Corsi Test and the control group’s Letter Span results. In the tDCS group, these correlations were absent, potentially due to greater interindividual variability in response to tDCS (Borella et al., 2017; Bruno et al., 2024; Shaw & Hosseini, 2020; Traut et al., 2021; Zinke et al., 2014). Furthermore, the scoring methods used for the N-back and OSPAN might have contributed to the lack of observed correlations, as these methods may lack the sensitivity needed to detect nuanced relationships between training and transfer measures (Conway et al., 2005; Tusch et al., 2016). For example, in the N-back, participants had a 50% chance of answering a trial correctly, whereas in the OSPAN, scoring was all-or-nothing (missing one number resulted in a score of zero).

### Neurophysiological Outcomes

The neurophysiological findings of this study revealed no significant differences between groups in oscillatory changes from pre- to post-training sessions. This suggests that tDCS combined with WM training was no more effective than WM training alone in eliciting oscillatory activity changes. These results align with our behavioural results and prior research (Chan et al., 2021b; Gordon et al., 2018; Šimko et al., 2021). However, significant differences were observed within both stimulation groups when comparing oscillatory activity across sessions, particularly in the changes between task phases rather than absolute power.

In the retention condition of the Letter Span, the control group showed a stronger increase in theta and high alpha power from the encoding to the retention phase during the post-training session compared to the pre-training session. In contrast, the tDCS group demonstrated no significant changes. These results may indicate that, despite stronger neural activity in the retention phase, the lack of a corresponding performance improvement could reflect a frustrated effort to enhance the capacity to protect stored information (Blacker et al., 2017; Langer et al., 2013; Manza et al., 2014; Michels et al., 2008; Roux & Uhlhaas, 2014). Additionally, the significant differences within the control group but not the tDCS group may highlight individual differences in neural adaptations, even after controlling for variables such as age, education, and baseline performance (Borella et al., 2017; Hsu et al., 2016; Shaw & Hosseini, 2020). These inconsistencies could explain why no significant correlations emerged between neural and behavioural changes.

In the manipulation condition of the Letter Span, both groups exhibited increased low alpha power from encoding to manipulation phases in the post-training session. The tDCS group also showed increased high alpha power from manipulation to recall phases. These changes suggest not only enhanced neural mechanisms for protecting stored information but also for manipulating it, which may have been a crucial factor in determining the effectiveness of the intervention in inducing significant improvements in WM performance (Blacker et al., 2017; Bonnefond & Jensen, 2012; Manza et al., 2014). Furthermore, another factor that may have contributed to the success of this compensatory mechanism is the greater cognitive demand of the manipulation condition compared to the retention condition. Retaining elements in memory while simultaneously reordering them was, according to our findings, significantly more challenging than merely retaining them. Nevertheless, the lack of significant neural-behavioural correlations and differences in the cortical regions associated with alpha power changes between groups warrant caution.

In the retention condition of the Corsi Test, consistent with the behavioural outcomes, no significant oscillatory changes were observed. However, in the manipulation condition, the control group demonstrated a reduction in theta power from the manipulation to the recall phase in the post-training session compared to the pre-training session. The pronounced decline in theta power in the left hemisphere may suggest two potential mechanisms underlying the outcomes. Firstly, it could indicate increased neural efficiency, with fewer frontal resources required during the recall phase to remember elements in the correct order (Cabeza et al., 2018; Duda & Sweet, 2020; Festini et al., 2018; Reuter-Lorenz & Cappell, 2008). Alternatively, it may reflect a reduction in hemispheric asymmetry induced by the training, a pattern typically observed in younger individuals and consistent with the HAROLD model (Cabeza et al., 2002, 2018; Erickson et al., 2007). We consider the latter explanation to be more plausible, as post-training scores on this test remained significantly lower compared to other tests. Importantly, the absence of comparable changes in the tDCS group may reflect variability in compensatory mechanisms or individual neural adaptations (Festini et al., 2018; Reuter-Lorenz & Park, 2014). Nonetheless, these interpretations remain speculative due to inconsistencies between groups, interindividual variability, and the lack of significant correlations.

The absence of robust tDCS effects may largely reflect individual differences in brain anatomy and neural plasticity, particularly in older adults (Krause & Cohen Kadosh, 2014; Polanía et al., 2018). Variability in cortical thickness, synaptic density, and baseline neural efficiency likely influences individual responsiveness to tDCS. Such diversity highlights the limitations of a one-size-fits-all approach and underscores the importance of personalised stimulation parameters (Kim et al., 2014; Opitz et al., 2015). For instance, certain individuals may require higher current intensities, longer stimulation durations, or alternative electrode montages to elicit optimal effects.

Personalising tDCS protocols could maximise its potential benefits but presents practical challenges. Techniques like HD-tDCS, which enhance current focality through multiple cathodal electrodes (Kuo et al., 2013; Parlikar et al., 2021), show promise but require further validation (Müller et al., 2022). Additionally, tailoring stimulation parameters may necessitate advanced neuroimaging tools like MRI and neuronavigation, increasing costs and logistical barriers, particularly in large-scale or clinical settings.

### Implications, Limitations, and Future Directions

Our findings align with meta-analyses that describe tDCS as “promising” but not consistently effective for WM enhancement in older adults (Goldthorpe et al., 2020; Indahlastari et al., 2021; Siegert et al., 2021; Vergallito et al., 2022). The findings of this study provide meaningful insights into the potential and limitations of tDCS as an adjunct to WM training in older adults. A key implication is the necessity of refining tDCS protocols to accommodate the unique anatomical and functional characteristics of older populations. Personalising tDCS parameters—such as stimulation intensity, duration, and electrode montage—may hold the key to improving its efficacy. This approach is especially pertinent given the substantial anatomical variability in the ageing brain, which undermines the traditional one-size-fits-all paradigm. Simply homogenising samples to reduce variability will likely be insufficient, as these anatomical differences can affect the distribution and intensity of the current, influencing individual responsiveness to tDCS.

Although no significant benefits of tDCS were observed in this study, our results underscore the importance of employing robust statistical approaches, such as LMEM, to address interindividual variability. By accounting for these differences, LMEM provided insights into patterns that may otherwise have been obscured. Additionally, the findings emphasise the critical role of neuroimaging tools in future research. EEG, for instance, not only complements behavioural measures but also reveals the neural dynamics underlying intervention effects. This study demonstrated the potential of adapted cognitive tasks to pinpoint specific phases where cognitive improvements occur, highlighting their value in advancing understanding of WM processes and intervention mechanisms.

Our results also support the notion that even short-term WM training can produce neuroplastic changes and cognitive benefits in older adults. However, the observed improvements were task-specific, suggesting that the selection of training tasks plays a crucial role in determining outcomes. Future intervention designs should prioritise tasks that engage multiple WM components to maximise training benefits and transfer effects.

Despite these implications, the study has several limitations. First, while the adapted tests were tailored to the study’s objectives, they may complicate comparisons with other research. Second, although groups were matched on demographic and baseline characteristics, the substantial contribution of random factors in our statistical models may have masked potential neural effects, particularly in the tDCS group. The study’s deliberate inclusion of a heterogeneous sample highlights the importance of examining individual differences but may have introduced variability that could not be fully controlled.

Third, while our spectral analyses of oscillatory activity provided valuable insights, we did not examine connectivity measures, which have been highlighted in prior studies as crucial for understanding the broader effects of tDCS on neural networks. Additionally, the spatial resolution limitations of EEG and the reliance on cluster-based permutation testing necessitate caution in interpreting the findings related to specific brain regions.

Looking forward, research on tDCS in older adults should focus on personalising stimulation protocols and integrating advanced neuroimaging tools to better understand its neural mechanisms. Key questions for future exploration include: How much current is required to induce meaningful changes? What is the optimal duration and intensity of stimulation? How durable are the observed effects? Addressing these questions will likely require combining approaches such as high-density tDCS, connectivity analysis, and longitudinal designs to explore the cumulative effects of training and stimulation.

By addressing these challenges, future studies can better assess the true potential of tDCS in enhancing WM and support its refinement into a more effective tool for cognitive interventions in older populations.

## CONCLUSION

In conclusion, this study found no significant additional benefits of tDCS in WM training for older adults. However, the WM training programme successfully induced neuroplastic changes and resulted in near-transfer and long-lasting effects, in both verbal and visuospatial WM tests, albeit only in the manipulation conditions. Notably, the WM training enhanced participants’ ability to retain and reorder information effectively. The underlying neural mechanisms differed between the Letter Span and Corsi Test: the former demonstrated greater increases in alpha power during the manipulation phase post-training, potentially indicating more effective efforts to maintain information while manipulating it, while the latter showed a marked reduction in left-hemisphere theta power during the recall phase post-training, possibly reflecting reduced hemispheric asymmetry.

Importantly, our findings were likely influenced by interindividual variability within the groups, which may have contributed to inconsistencies between groups and the lack of supporting correlations in neural outcomes. This suggests caution in interpreting these results. Nonetheless, these findings underscore the importance of personalising tDCS parameters to uncover its true effects, emphasising the need for neuroimaging tools to analyse tDCS outcomes more effectively. Additionally, these results support a more optimistic view of adaptive WM training, indicating that it can deliver lasting behavioural and neuroplastic benefits in older adults after just a few sessions.

## SUPPLEMENTARY MATERIAL

**Figure 7.**
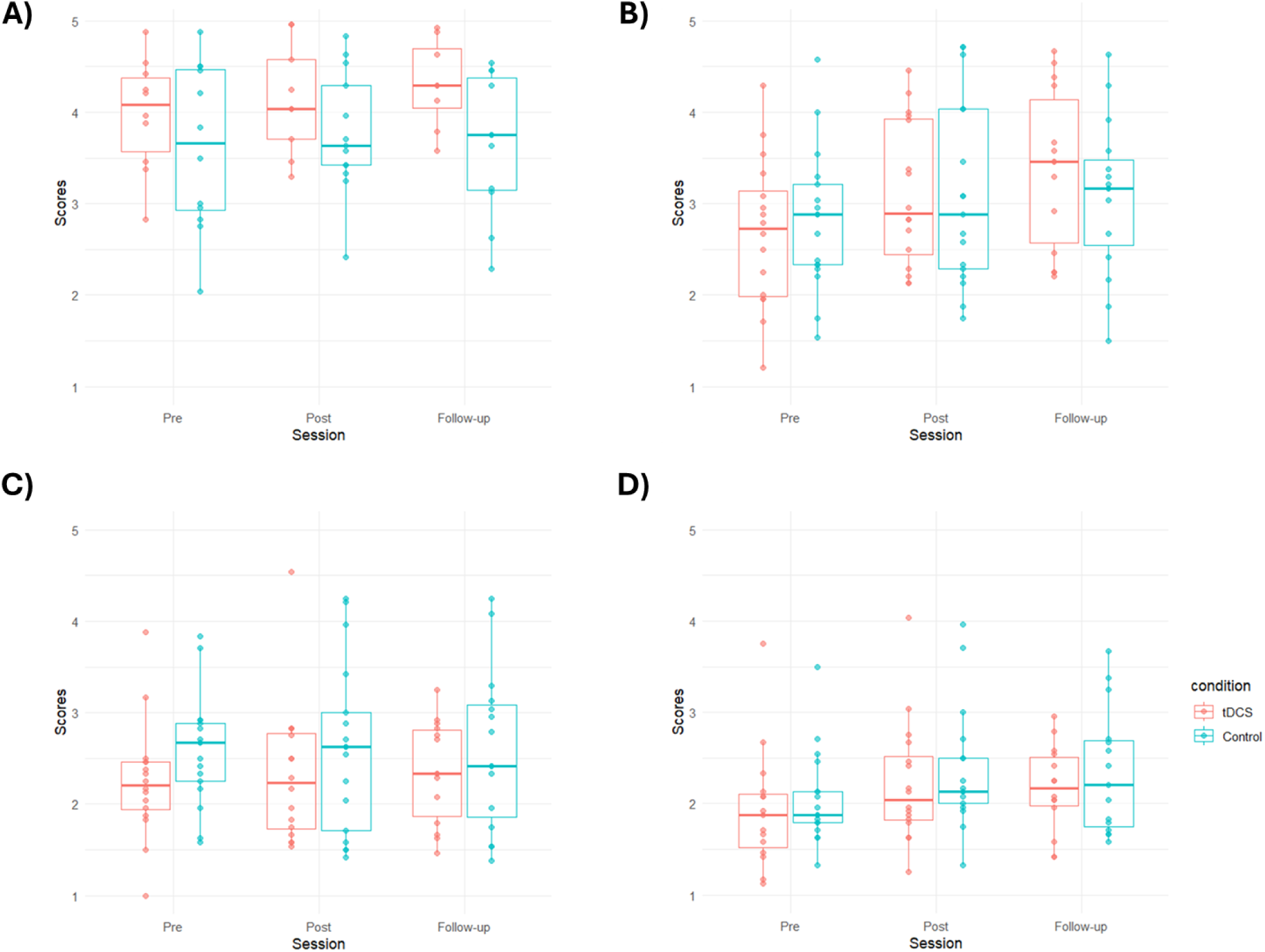
Mean scores and confidence intervals of the retention and manipulation conditions of the Letter Span and the Corsi Test. A) Retention – Letter Span. B) Manipulation – Letter Span. C) Retention – Corsi Test. D) Manipulation – Corsi Test.

